# Whale shark rhodopsin adapted to its vertically wide-ranging lifestyle

**DOI:** 10.1101/2021.10.01.462724

**Authors:** Kazuaki Yamaguchi, Mitsumasa Koyanagi, Keiichi Sato, Akihisa Terakita, Shigehiro Kuraku

**Affiliations:** Laboratory for Phyloinformatics, RIKEN Center for Biosystems Dynamics Research (BDR), Kobe, Japan; Department of Biology and Geosciences, Graduate School of Science, Osaka City University, Osaka, Japan; Okinawa Churashima Research Center, Okinawa Churashima Foundation, Okinawa, Japan; Okinawa Churaumi Aquarium, Okinawa, Japan; Molecular Life History Laboratory, Department of Genomics and Evolutionary Biology, National Institute of Genetics, Mishima, Japan; Department of Genetics, Sokendai (Graduate University for Advanced Studies), Mishima, Japan

**Keywords:** visual opsin, rhodopsin, whale shark, spectral tuning, night blindness, thermal stability

## Abstract

Spectral tuning of visual pigments often facilitates adaptation to new environments, and it is intriguing to study the visual ecology of pelagic sharks with expanded habitats. The whale shark, which dives into the deep sea of nearly 2,000 meters besides near-surface filter-feeding, was previously shown to possesses the ‘blue-shifted’ rhodopsin (RHO). In this study, our spectroscopy of recombinant whale shark RHO mutants revealed the dominant effect of the novel spectral tuning amino acid site 94, which is implicated in congenital stationary night blindness of humans, accounting for the blue shift. Thermal decay profiling revealed the reduction of the thermal stability of whale shark RHO, as typically observed for cone opsins, which was experimentally shown to be achieved by the site 178, as well as 94. The results suggest that these two sites cooperatively enhance the visual capacity in both the deep sea and the sea surface, enabling exceptionally wide vertical migration of this species.

## Introduction

The whale shark *Rhincodon typus*, known as the largest extant fish, dives into the deep sea of 2,000 meters while it forages with filter-feeding near the surface (Tyminski et al., 2015). Because of its exceptionally wide vertical habitat range, it is of great interest to investigate its visual ecology. Recent genome sequencing confirmed degenerated visual opsin gene repertoires of this species, containing only rhodopsin (RHO or Rh1) and long wavelength-sensitive opsin (LWS) (Hara et al., 2018). Our previous spectroscopic analysis on the whale shark RHO pigment, performed *in vitro*, uncovered a remarkable shift of the wavelength of the maximum absorbance (λmax) to 478 nm, from the presumable ancestral condition of 500 nm (Figure 1; Hara et al., 2018). As the light of approximately 480 nm is the least attenuated in the deep sea, this finding suggested the reliance of the whale shark on vision in the deep sea with the tuned RHO, but the mechanism that allows this so-called ‘blue shift’ has remained unknown. Interestingly, an *in silico* study on the whale shark RHO has failed to predict candidate amino acid residues that account for the blue shift (Fasick et al., 2019).

**Figure 1.**
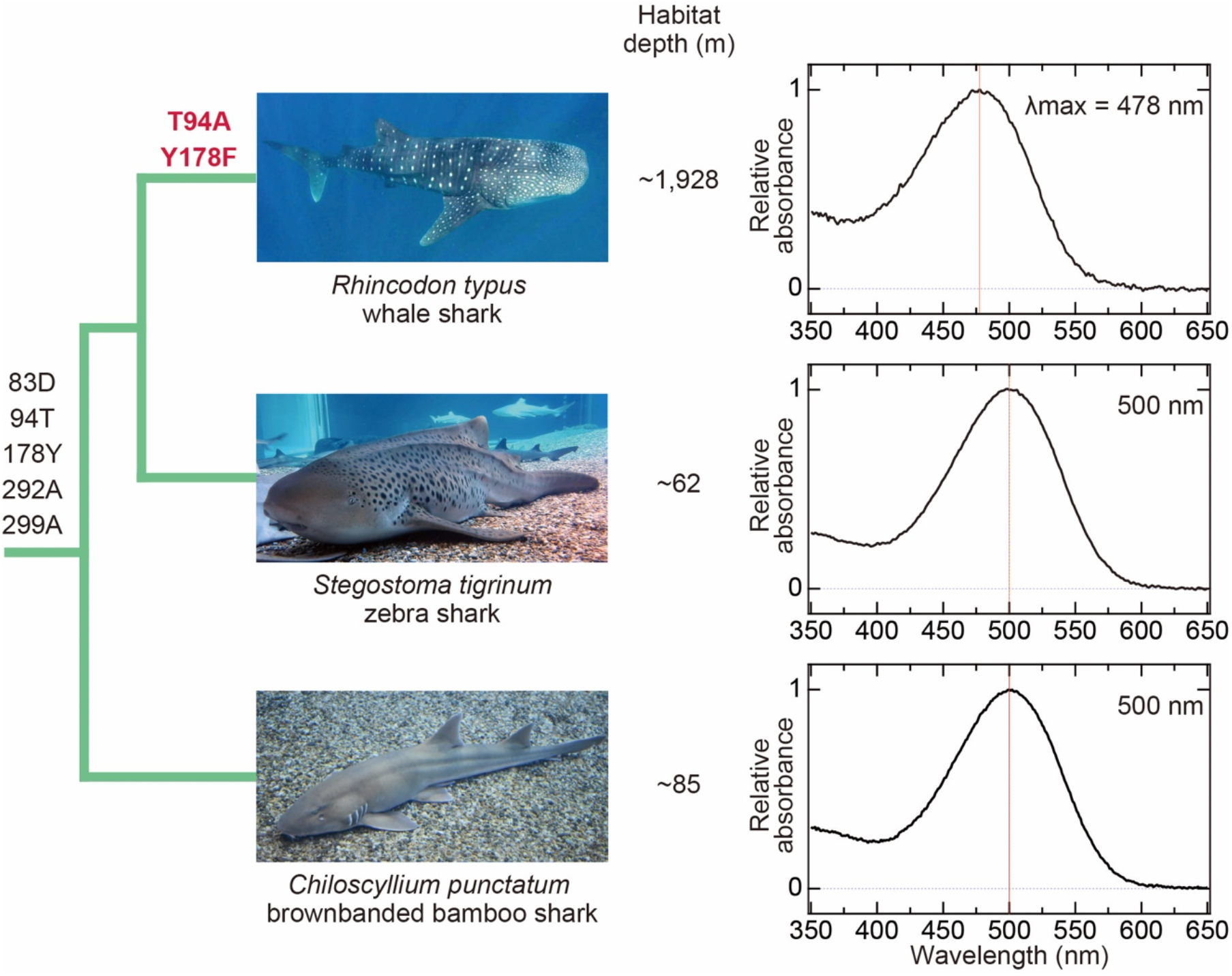
Absorption spectra of RHO pigments of the whale shark and its close relatives. Their phylogenetic relationship is depicted with the amino acid residues at novel (94 and 178) and previously documented (83, 292, and 299) spectral tuning sites. Each absorption spectrum was normalized to the maximum absorbance in the visible light region. The wavelengths of maximum absorbance of the RHO pigment (λmax) are based on this study (zebra shark) and existing literature (Hara et al., 2018).

## Results

We first focused on the zebra shark *Stegostoma tigrinum* (formerly, called *S. fasciatum*), one of the phylogenetically closest extant species to the whale shark (Figure 1; Naylor et al., 2012). The absorption spectrum of the zebra shark RHO peaked at 500 nm, as in the other close relative dwelling in shallow waters, the brownbanded bamboo shark (Figure 1; Hara et al., 2018). This substantiated the uniqueness of the blue shift to the whale shark lineage.

Next, we scrutinized the sequence alignment of RHO orthologs of these sharks and their relatives. This analysis did not reveal any amino acid substitution unique to the whale shark among the previously characterized ‘spectral tuning sites’ of the RHO, such as the sites 83, 292, and 299 (Hart et al., 2020; Musilova et al., 2019; Yamaguchi et al., 2021; Figure 1—figure supplement 1). Among the amino acid residues positioned within the range of 4.5 Å from the chromophore 11-*cis*-retinal (Palczewski et al., 2000), we identified two residues 94 and 178 substituted exclusively in the whale shark RHO (Ala94 and Phe178; Figure 1—figure supplement 1). We examined the effect of each of these substitutions with site-directed mutagenesis. Spectroscopic analysis of recombinant RHO mutants revealed that the λmax of whale shark RHO A94T and F178Y mutants shifted toward longer wavelengths by 19 and 3 nm, respectively, compared with that of the wild-type (Figure 2A). Conversely, mutants with substitutions at these sites (T94A and Y178F) of zebra shark recapitulated the blue shift by 19 and 3 nm (Figure 2B). This result shows a dominant effect of Ala94 in the blue shift of the whale shark RHO and accounts for the 22 nm difference from the λmax of zebra shark RHO by cumulative effect of these two mutations. At the site 94 of RHO, no case of natural substitution has been reported as responsible for spectral tuning, except for the T94I causing a certain type of human diseases including congenital stationary night blindness (CSNB) (Al-Jandal et al., 1999; reviewed in Park, 2014). The molecular basis of CSNB with T94I is explained by the substitution from a hydrophilic to hydrophobic residue, which leads to lower thermal stability via thermal isomerization of the retinal and hydrolysis of the Schiff base (Singhal et al., 2016).

**Figure 2.**
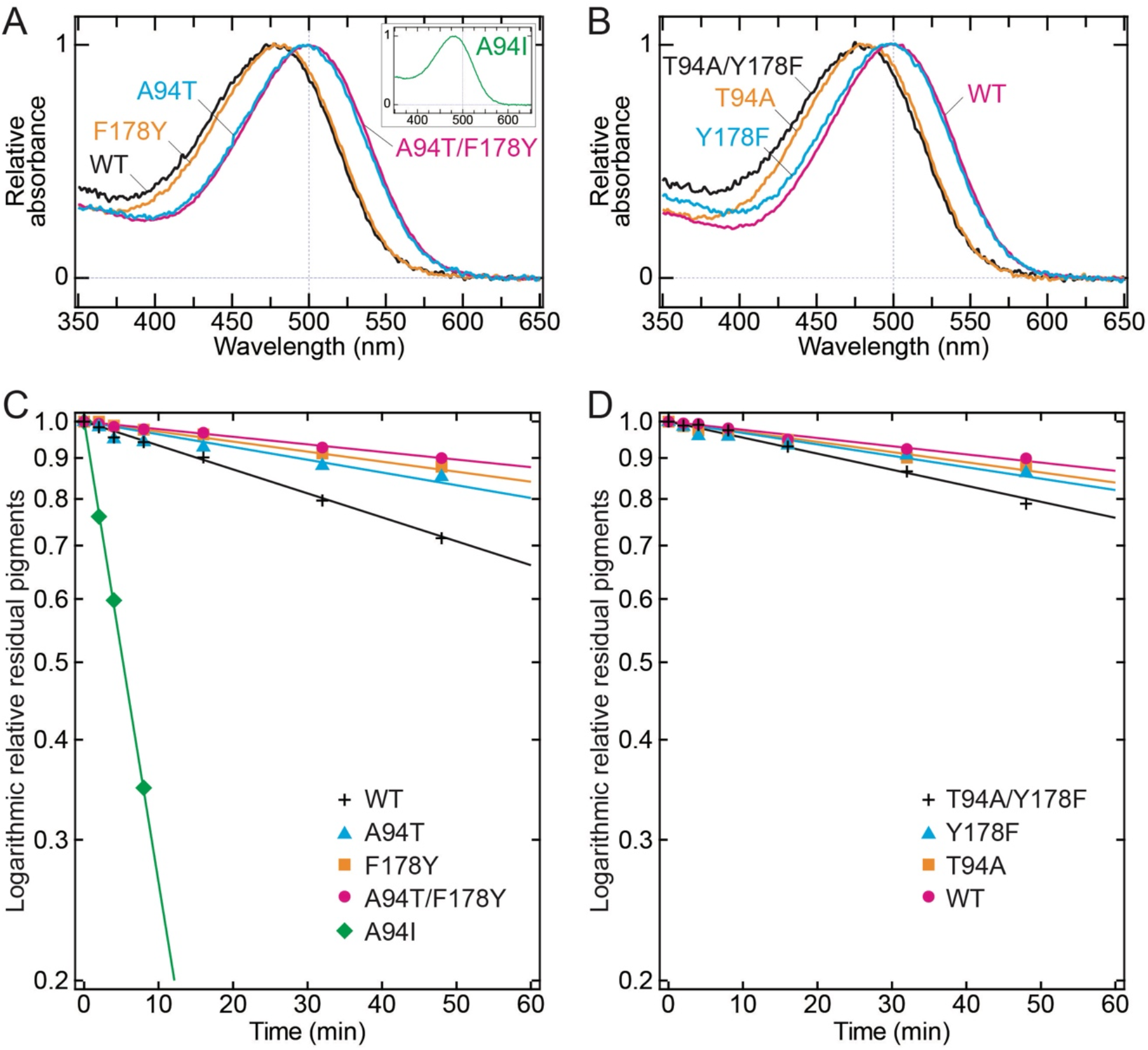
Experimental characterization of the whale shark and the zebra shark RHOs. (A) Absorption spectra of the wild-type (WT, black), the A94T mutant (blue), the F178Y mutant (orange), the A94T/F178Y mutant (magenta), and the A94I mutant (green; inset) of the whale shark RHO. (B) Absorption spectrum of the wild-type (WT; magenta), the T94A mutant (orange), the Y178F mutant (blue), and the T94A/Y178A mutant (black) of the zebra shark RHO. (C) Time-course thermal decay for the whale shark RHO. The colors correspond to those for the constructs in A. The vertical axis shows relative amounts of residual RHO quantified with the average absorbance of individual RHO samples at the 5 points (λmax-2, λmax-1, λmax, λmax+1, and λmax+2 nm), assuming that the absorbance immediately after heating was 1 and the absorbance after light irradiation was 0. (D) Time-course thermal decay for the zebra shark RHO. The colors correspond to those for the constructs in B, and the vertical axis is shown in the same way as in C.

This observation in humans prompted us to investigate thermal stability of the whale shark RHO (Figure 2C, D). Similar to the case of T94I included above (Singhal et al., 2016), the A94I mutant of the whale shark RHO rapidly decayed at 37 °C, indicating that the site 94 also affects the thermal stability in the whale shark RHO (Table 1 and Figure 2C). In fact, the wild-type whale shark RHO (Ala94) also showed faster decay with a half-life (t_1/2_) of 99.0 min than its A94T mutant (t_1/2_= 173.3 min) (Figure 2C—figure supplement 2, Table 1), indicating a decrease of thermal stability by the T94A substitution. Interestingly, the whale shark RHO A94T mutant has higher thermal stability than the wild-type but still lower than the zebra shark RHO wild-type (Thr94, t_1/2_ = 346.6 min). This result is consistent with the observation that the zebra shark RHO T94A mutant has higher thermal stability (t_1/2_ = 231.0 min) than the whale shark RHO wild-type (Ala94, t_1/2_ = 99.0 min) (Figure 2C, D).

**Table 1.** Spectral and thermal properties of the wild-type and mutant RHOs studied.

| Rhodopsin construct | Amino acid residue at site 94 | Amino acid residue at site 178 | $\lambda_{\text{max}}$ (nm) | Difference from wild-type (nm) | Estimated half-life of rhodopsin* (min) |
| --- | --- | --- | --- | --- | --- |
| whale shark wild-type | A | F | 478 | NA | 99.0 |
| whale shark A94T | T | F | 497 | +19 | 173.3 |
| whale shark A94I | I | F | 478 | 0 | 5.3 |
| whale shark F178Y | A | Y | 481 | +3 | 231.0 |
| whale shark A94T/F178Y | T | Y | 500 | +22 | 346.6 |
| zebra shark wild-type | T | Y | 500 | NA | 346.6 |
| zebra shark T94A | A | Y | 481 | -19 | 231.0 |
| zebra shark Y178F | T | F | 497 | -3 | 231.0 |
| zebra shark T94A/Y178F | A | F | 478 | -22 | 138.6 |

\*Time duration for the relative absorbance of rhodopsin to reach 0.5. The exponential trendlines for determining the half-life values ( $t_{1/2}$ ) were calculated from the relative absorbance plots of RHOs using all measured data points. NA, not applicable.

We also scrutinized the site 178, the other site substituted exclusively in the whale shark RHO. The F178Y mutant of the whale shark RHO exhibited comparable thermal stability (t_1/2_= 231.0 min) to the A94T mutant, and introducing both A94T and F178Y substitutions into the whale shark RHO recovered high thermal stability (t_1/2_ = 346.6 min) close to that of the zebra shark wild-type (Figure 2C). A parallel effect was observed in the zebra shark RHO; mutations at both sites (T94A/Y178F) resulted in a severe decrease of thermal stability than either mutation (Figure 2D). Taken together, substitutions at the two sites and their cumulative effect account for the lower thermal stability of the whale shark RHO (Table 1), as in the blue shift. The rate of half-life difference between the wild-type and the double mutant for the whale shark (≈3.5 times) exceeded that for the zebra shark (≈2.5 times) (Table 1), which could be explained by possible influence from other amino acid substitutions that accumulated in either evolutionary lineage.

## Discussion

Blue shift of RHO resulting from deep-sea life has repeatedly been documented, but mostly by studies on deep-sea ‘specialists’ (Hart et al., 2020; Musilova et al., 2019). In contrast, the whale shark is reported to spend a considerable proportion of the daytime on filter-feeding near the surface, besides occasional deep dives down to nearly 2,000 meters (Tyminski et al., 2015). Although the whale shark is capable of stabilizing deep body temperatures (Nakamura et al., 2020), the retina is thought to be affected by wide-ranging water temperatures that depend on the depth (e.g., 4.2 to 33.0 °C reported in Tyminski et al., 2015). Our present study revealed novel spectral tuning sites for blue shift of RHO, Ala94 and Phe178, which could have enabled its unique lifestyle with exceptionally wide vertical migration.

Our study revealed a commonality of the substitution at the site 94 in the blue-shifted spectrum and lowered thermal stability between the whale shark RHO and the human RHO with a CSNB-causing mutation (T94I). In general, high thermal stability is a critical feature of RHO to achieve a high signal-to-noise ratio by reducing dark noise, which inherently enables dim-light vision (Yanagawa et al., 2015). This functional nature of RHO constraining dark noise usually does not permit the sites 94 or 178 to contribute to its spectral tuning. The co-existence of these substitutions has not been observed in any visual opsins orthologous to RHO by our exhaustive search in the public sequence database (Supplementary Tables 1 and 2).

In our experiment at 37 °C, the thermal stability of whale shark RHO excelled that of the T94I mutant (Figure 2C and Table 1—figure supplement 2) but is significantly lower than that of zebra shark (Figure 2C, D). The reduced thermal stability of the whale shark RHO is reminiscent of cone opsins that have lower thermal stability than RHOs to enable daylight vision. In fact, compared to the human RHO (Janz et al., 2003), the whale shark RHO has lower thermal stability, which is rather closer to that of mammalian LWS/MWS (Alexander et al., 2017; Srinivasan et al., 2017) and chicken RH2 (Sato et al., 2018). Near the sea surface, the cone opsin-like low thermal stability of whale shark RHO in high-temperature water (Figure 2C) can facilitate the photoreceptor light adaptation suitable for the bright light condition. While the whale shark has evolutionarily lost all cone opsins except LWS (Hara et al., 2018), the whale shark RHO may compensate the degenerative cone opsin repertoires by facilitating light adaptation. On the other hand, in the deep sea, the functionality of whale shark RHO to reduce dark noise would be maintained at low temperatures, as shown for RHO mutants with persistent functionality at low temperatures (Janz et al., 2003). This speculation is compatible with the blue shift of the whale shark RHO that is thought to facilitate dim-light sensing in the deep sea.

Our *in vitro* study that is supported by cross-species, genome-scale sequence informatics provides the first case of spectral tuning of RHO by natural substitutions at the sites 94 and 178. The former plays a major role in the blue shift analyzed in this study, while the pivotal role of the latter is regulating thermal stability. The route of “rhodopsin tuning” we revealed is distinct from typical solutions for deep-sea specialists and may have facilitated the habitat expansion of this iconic species with vertically wide movement utilizing both deep sea and sea surface.

## Materials and Methods

### cDNA sequencing

The eyes of a female zebra shark that was born from captive breeding were sampled at the Okinawa Churaumi Aquarium in accordance with the official guideline for animal experiments at Okinawa Churaumi Aquarium. Total RNAs were extracted from these tissues using TRIzol Reagent (Life Technologies) and used as a template in reverse transcription with the SMART cDNA Library Construction Kit (Clontech). cDNA fragments amplified using the degenerate primers designed based on the ortholog sequences of close relatives were sequenced with a 3730xl DNA Analyzer (Thermo Fisher Scientific). The nucleotide sequence of the zebra shark RHO is deposited in GenBank under the accession ID MT625929. Our experiment was conducted in accordance with the institutional guideline Regulations for the Animal Experiments and approved by the Institutional Animal Care and Use Committee (IACUC) of RIKEN Kobe Branch.

### Rhodopsin expression and purification

The RHO expression vectors designed as described previously (Koyanagi et al., 2015) based on the abovementioned zebra shark sequence as well as the previously reported whale shark sequence (Accession ID MT625928; Hara et al., 2018) were synthesized and cloned by a custom service at GenScript. The RHO mutants were constructed using the KOD-Plus-Mutagenesis Kit (Toyobo). The vectors were transfected into HEK293S cells by the calcium-phosphate method. The cells were cultured for 48 hours for protein expression. The RHO pigments were reconstituted as described previously (Hara et al., 2018). An excessive amount of 11-*cis*-retinal was added to the cells recovered by centrifugation, and the pigments were constituted by overnight incubation at 4 °C in the dark. The cells were lysed in 1% (w/v) n-dodecyl-β-D-maltoside (DM) in 50 mM HEPES buffer (pH 6.5) containing 140 mM NaCl (Buffer P), and the pigments were solubilized. The pigments were bound to 1D4-agarose, washed with 0.02% (w/v) DM in Buffer P, and eluted with 0.02% (w/v) DM in Buffer P containing the 1D4 peptide.

### Spectroscopic measurements

Absorption spectra of purified pigments were measured with the V-750 UV-VIS Spectrophotometer (JASCO International). Yellow light was supplied by the light source with Y-50 glass cutoff filter (Toshiba). The maximum absorption spectra were measured at 4 °C in the dark. Thermal decay assay of RHO was performed with incubation at 37 °C in the dark, and the λmax was measured every two minutes. At the end of the measurement, the pigments were irradiated with light, and the spectra of the bleached pigments were measured.

### *In silico* scan of RHO sequences

The NCBI nr database as of June 2020 was downloaded for a BLASTP search using the whale shark RHO sequence as a query. The collected peptide sequences were aligned with the program MAFFT v7.299b using the L-INS-i method (Katoh & Standley, 2013). The aligned sequences were trimmed with trimAl v1.4.rev15 using the ‘-automated1’ option, followed by the removal of gapped sites using the ‘-nogaps’ option (Capella-Gutiérrez et al., 2009). The maximum-likelihood tree was inferred with RAxML v8.2.8 using the PROTCATWAG model. For evaluating the confidence of nodes, the rapid bootstrap resampling with 1,000 replicates was performed (Stamatakis, 2014). Based on the inferred phylogenetic tree, 6,803 sequences regarded as RHO orthologs were extracted for a scan of the amino acid residues at the sites 94 and 178.

## Acknowledgments

We thank Rui Matsumoto and Kiyomi Murakumo at Okinawa Churaumi Aquarium for their assistance in sampling. Our gratitude extends to Tomohiro Sugihara at Osaka City University for his assistance in laboratory experiments.

## Additional information

### Author contributions

S.K. and A.T. designed the study. K.Y., K.S. and M.K. performed research and analyzed the data. S.K., K.Y., and M.K. drafted the manuscript, and all the authors contributed to manuscript finalization.

