## Supplementary Information for "Whale shark rhodopsin adapted to its vertically wide-ranging lifestyle"

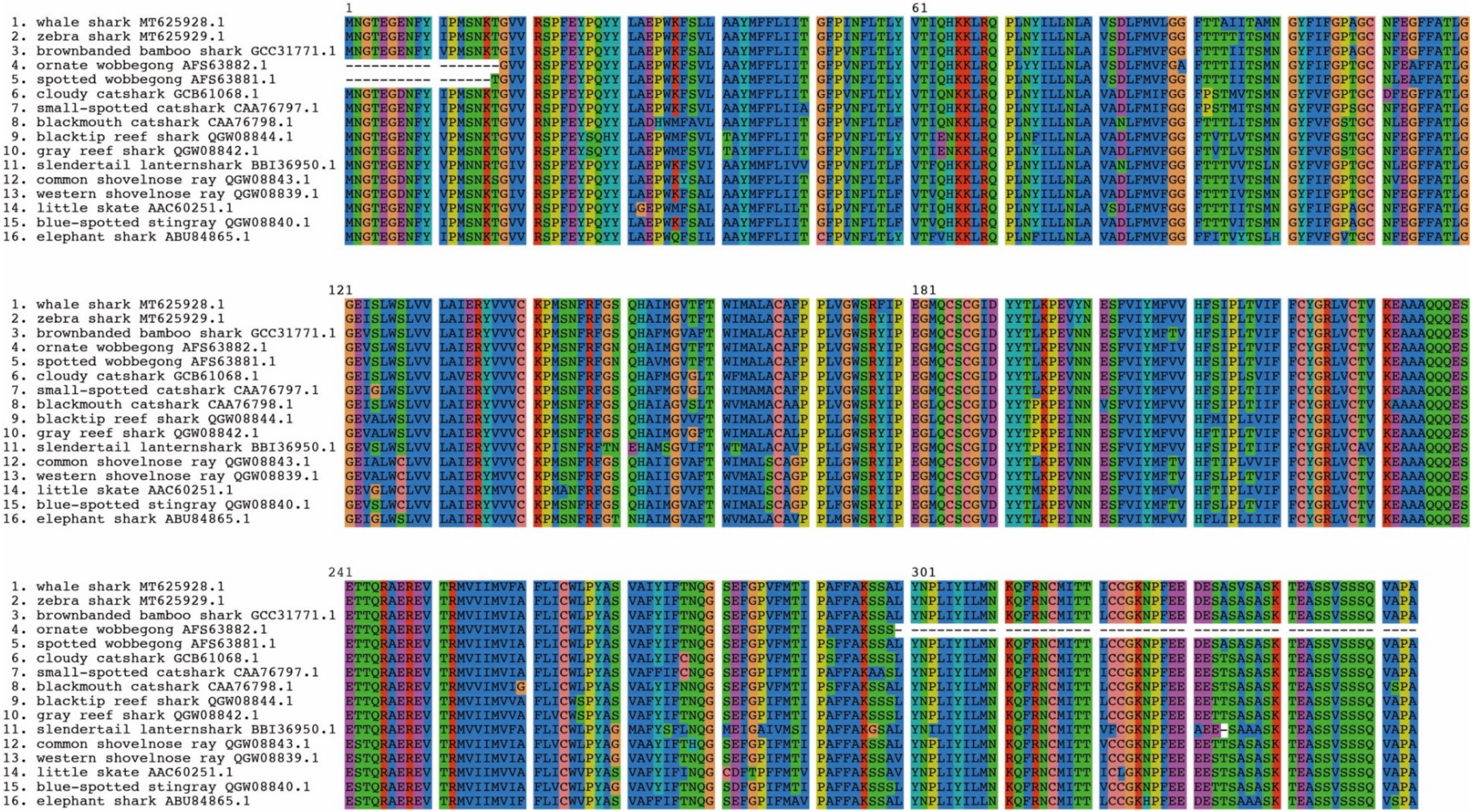


figure supplement 1. Multiple alignment of amino acid sequences of cartilaginous fish RHOs. Hyphens indicate gaps or undetermined regions because of incomplete sequencing.

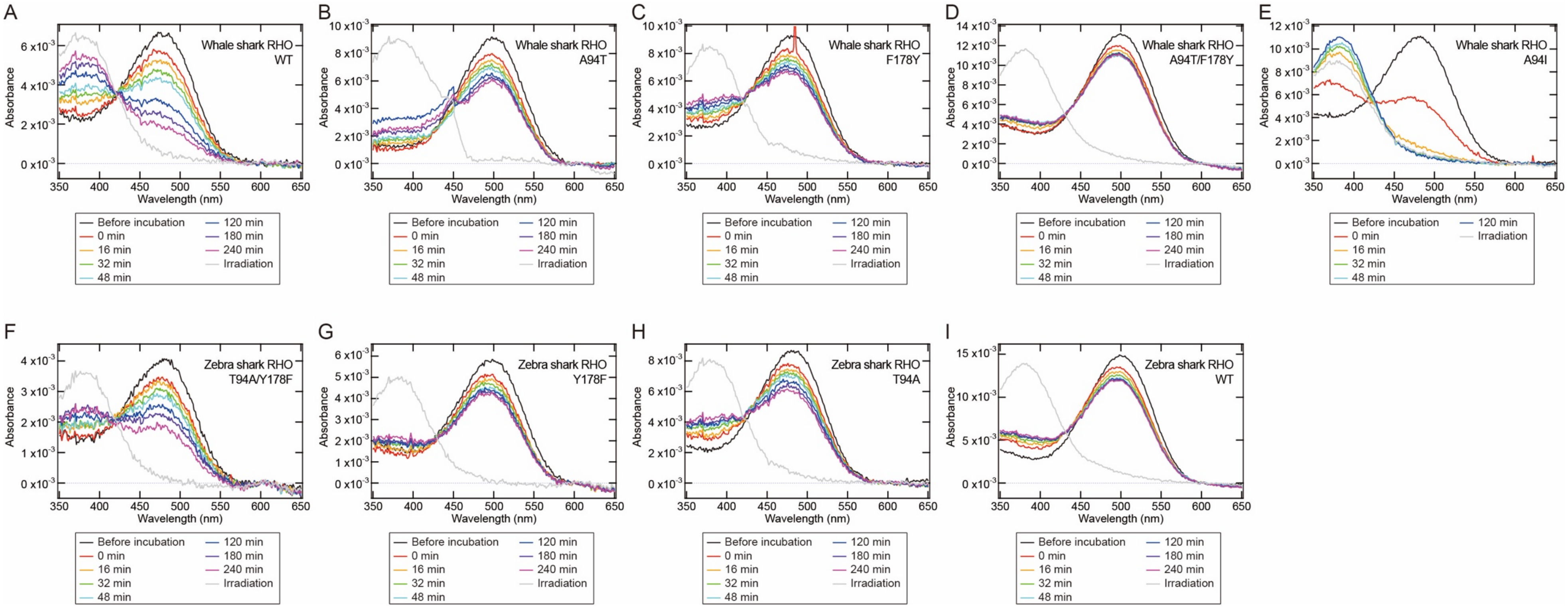


figure supplement 2. Absorption spectra of RHOs under 37 °C heating conditions in time course. (A) whale shark RHO wild-type (WT). (B) whale shark RHO A94T. (C) whale shark RHO F178Y. (D) whale shark RHO A94T/F178Y. (E) whale shark RHO A94I. (F) zebra shark RHO T94A/Y178F. (G) zebra shark Y178F. (H) zebra shark T94A. (I) zebra shark RHO WT.

Supplementary Table 1. Teleost fish species with substitution at site 94 of RHO.

| Species name | Ortholog group name | Amino acid residue at site 94 | Accession ID | Habitat# |
|---|---|---|---|---|
| Antarctic toothfish (*Dissostichus mawsoni*) | Exo-rhodopsin | A | KAF3860639.1 (JAAKFY010000002.1*) | habitat depth: 0 - 2,200 m (usually 88 - 1,600 m); latitude/longitude: 45°S - 78°S, 180°W - 180°E |
| emerald rockcod (*Trematomus bernacchii*) | Exo-rhodopsin | A | XP_033982177.1 | habitat depth: 0 - 700 m (usually 0 - 200 m); latitude/longitude: 61°S - 78°S |
| ploughfish (*Gymnodraco acuticeps*) | Exo-rhodopsin | A | XP_034066829.1 | habitat depth: 0 - 550 m; latitude/longitude: 61°S - 78°S |
| black rockcod (*Notothenia coriiceps*) | Exo-rhodopsin | A | XP_010794173.1 | habitat depth: 0 - 550 m; latitude/longitude: 46°S - 78°S, 180°W - 180°E |

*The putative open reading frame was manually estimated by us from the genomic sequence because the open reading frame registered in GenBank was incorrect. #Based on the FishBase (https://www.fishbase.se/).

Supplementary Table 2. Species with substitution at site 178 of RHO.

| Species name | Ortholog group name | Amino acid residue at site 178 | Accession ID | Habitat[#] |
|---|---|---|---|---|
| Gaboon caecilian (*Geotrypetes seraphini*) | rhodopsin | F | XP_033782390.1 | lives mainly underground in lowland forest |
| two-lined caecilian (*Rhinatrema bivittatum*) | rhodopsin | F | XP_029453475.1 | not available |
| black shield tail snake (*Melanophidium khairei*) | rhodopsin | F | AOF40379.1 | latitude/longitude: 15° 57' N, 73° 59' E; elevation: 715 m |
| wolf eel (*Anarrhichthys ocellatus*) | rhodopsin | F | XP_031700636.1 | habitat depth: 1 - 226 m; latitude/longitude: 80°N - 26°N, 118°E - 111°W |
| pirate perch (*Aphredoderus sayanus*) | rhodopsin | F | AAZ23943.1 | temperate: 5°C - 26°C; latitude/longitude: 46°N - 28°N |
| *Astronesthes chrysophekadion* | rhodopsin | F | AHA33545.1 | habitat depth: 100 - 1,120 m |
| Mexican tetra (*Astyanax mexicanus*) | rhodopsin | F | XP_015463133.1 | latitude/longitude: 36°N - 24°N |
| *Astyanax ruberrimus* | rhodopsin | F | QJT41803.1 | not available |
| spiderfish (*Bathypterois dubius*) | rhodopsin | F | AAN08922.1 | habitat depth: 750 - 1,941 m; temperate: 4°C - 12°C |
| swampfish (*Chologaster cornuta*) | rhodopsin | F | AEJ09054.1 | latitude/longitude: 38°N - 32°N; temperate: - 23°C; in cave |
| cave gudgeon (*Milyeringa veritas)* | rhodopsin | F | AGR53615.1 | habitat depth: 0 - 32 m; in cave |
| Alabama cavefish (*Speoplatyrhinus poulsoni*) | rhodopsin | F | AEJ09058.1 | latitude/longitude: 35°N - 34°N; in cave |
| grey cutthroat (*Synaphobranchus affinis*) | rhodopsin | F | AYW11990.1 | habitat depth: 290 - 2,400 m (usually 400 - 500 m); temperate: 3°C - 11°C |

| | | | | |
|---|---|---|---|---|
| *Typhlichthys eigenmanni* | rhodopsin | F | AEJ09077.1 | not available |
| southern cavefish (*Typhlichthys subterraneus*) | rhodopsin | F | AEJ09062.1 | latitude/longitude: 39°N - 34°N; in cave |
| *Sinocyclocheilus purpureus* | rhodopsin | C | ACF22658.1 | not available |
| pallid sculpin (*Cottunculus thomsonii*) | rhodopsin | H | AAR24239.1 | habitat depth: 100 - 1,600 m; latitude/longitude: 73°N - 10°N, 81°W - 2°W |

#Based on AmphibiaWeb (https://amphibiaweb.org/), The Reptile Database (https://reptile-database.reptarium.cz/), and FishBase (https://www.fishbase.se/).
